# Exploring the Categorical Nature of Colour Perception: Insights from Artificial Networks

**DOI:** 10.1101/2024.01.25.577209

**Authors:** Arash Akbarinia

**Affiliations:** Department of Experimental Psychology, Justus Liebig University Giessen, Germany

**Keywords:** colour categories, colour naming, colour perception, deep neural networks, artificial psychophysics

## Abstract

This study delves into the categorical aspects of colour perception, employing the odd-one-out paradigm on artificial neural networks. We reveal a significant alignment between human data and unimodal vision networks (e.g., ImageNet object recognition). Vision-language models (e.g., CLIP text-image matching) account for the remaining unexplained data even in non-linguistic experiments. These results suggest that categorical colour perception is a language-independent representation, albeit partly shaped by linguistic colour terms during its development. Exploring the ubiquity of colour categories in Taskonomy unimodal vision networks highlights the task-dependent nature of colour categories, predominantly in semantic and 3D tasks, with a notable absence in low-level tasks. To explain this difference, we analysed kernels’ responses before the winnertaking-all, observing that networks with mismatching colour categories align in continuous representations. Our findings quantify the dual influence of visual signals and linguistic factors in categorical colour perception, thereby formalising a harmonious reconciliation of the universal and relative debates.

## 1 Introduction

The electromagnetic spectrum of light reaching our eyes presents a seamless continuum, devoid of any apparent discontinuities. However, our visual system transforms this continuous spectrum into distinct colour categories, as exemplified by the captivating hues of the rainbow. This prompts a fundamental question: why does our perceptual system organise a continuous function into discrete colour categories? If this discretisation were merely a computational expedient, colour categories would be uniformly distributed. Yet, there is a large variation among the volume occupied by colour categories, for instance, green and blue dominate extensive segments, while yellow and brown occupy more confined spaces.

Numerous studies in the literature have delved into this phenomenon, proposing two competing theories. Universalists [6] argue that the mechanism underpinning categorical colour perception is an inherent aspect of physiological processes. Conversely, relativists [13] posit that language and culture play a pivotal role in shaping colour categories. Universalists bolster their argument by citing the overlap in focal colours across diverse cultures [33] and experiments utilising nonverbal paradigms [22]. Relativists highlight the challenges children encounter in acquiring colour names [35] and the variations in colour terms across languages [36]. Scientific consensus has oscillated between these perspectives, eventually settling on a compromise position of moderate universality: universal patterns beyond superficial discrepancies across different cultures (see the review by Kay and Regier [24]). Nonetheless, two open questions persist in this position. First, isolating the primary driving force behind the emergence of colour categories is unfeasible given the intricate interplay between linguistic and perceptual processing. Second, if the universalism theory is favoured, it is unclear why our visual system adopts a categorical colour representation—is this due to the neural circuitry of our system or linked to the visual tasks we perform?

This article addresses these inquiries by harnessing the capabilities of artificial neural networks (ANNs), which possess sufficient complexity to emulate the ecological validity of human observers while remaining amenable to controlled experiments. Previous studies have utilised unimodal ANNs to investigate colour categories. Chaabouni et al. [10] demonstrated that the accuracy-complexity trade-off in human colour terms emerges in two artificial agents playing a communication game. This finding aligns with efficient communication theory, which asserts that human colour categories closely approach the theoretically optimal limit [50, 20], thereby reinforcing the pivotal role of language in shaping colour categories. In a contrasting approach, de Vries et al. [14] illustrated that colour boundaries reported by human observers manifest in object recognition networks trained on natural images without any language component. This observation aligns with categorical perception theory, asserting that perceptual colour space is warped by stretching at category boundaries or by within-category compression [8, 48, 45]. Consequently, this finding suggests that colour categories may develop independently of language. In this study, we employ linear probes [5] to (1) compare multimodal vision-language and unimodal vision deep neural networks, thereby dissecting the contribution of each modality, and (2) scrutinise the representation in an identical architecture (ResNet50) trained on different visual tasks to explore whether the system’s functional role influences categorical colour representation.

Our investigation has yielded insightful findings. Firstly, we offer a resolution to the enduring debate between universalists and relativists. Unimodal vision models, exemplified by ImageNet object recognition networks, explain over eighty per cent of human data, leaving the remaining unexplained portion attributed to multimodal vision-language models, such as CLIP text-image matching networks. This underscores that categorical colour perception constitutes a language-independent representation, despite the discernible influence exerted by linguistic colour terms on its developmental trajectory. Secondly, our findings reveal that human-like colour categories predominantly emerge in models trained on semantic visual tasks, including image segmentation, object recognition, and scene classification. Networks optimised for 3D tasks exhibit moderately human-like colour categories, while those focused on 2D lowlevel tasks, such as autoencoding and denoising, fall short of reproducing human-like colour categories. Lastly, our investigation underscores that networks with distinct discrete colour categories may possess a highly similar underlying continuous representation of how colour is partitioned in space. However, following the process of discretisation (winner-take-all), the output categories do not align.

## 2 Results

We systematically investigated the categorical colour representation within artificial neural networks, utilising Munsell chips (see insert **d** in Fig. 1). This set gained prominence through its inclusion in the World Colour Survey (WCS) [23] and is frequently employed in colour category literature (e.g., [34, 29, 50, 10]). In our analysis, we compared the outputs of artificial networks with human data from [6, 41], concentrating on eight chromatic colour categories: red, orange, yellow, brown, green, blue, purple, and pink. To enhance the reliability of our findings, each Munsell chip was tested as the surface colour of 2904 superellipse shapes (see Fig. A1). To investigate the role of language and visual signals in categorical colour perception, we examined two types of networks: unimodal vision and multimodal language-vision models.

**Fig. 1.**
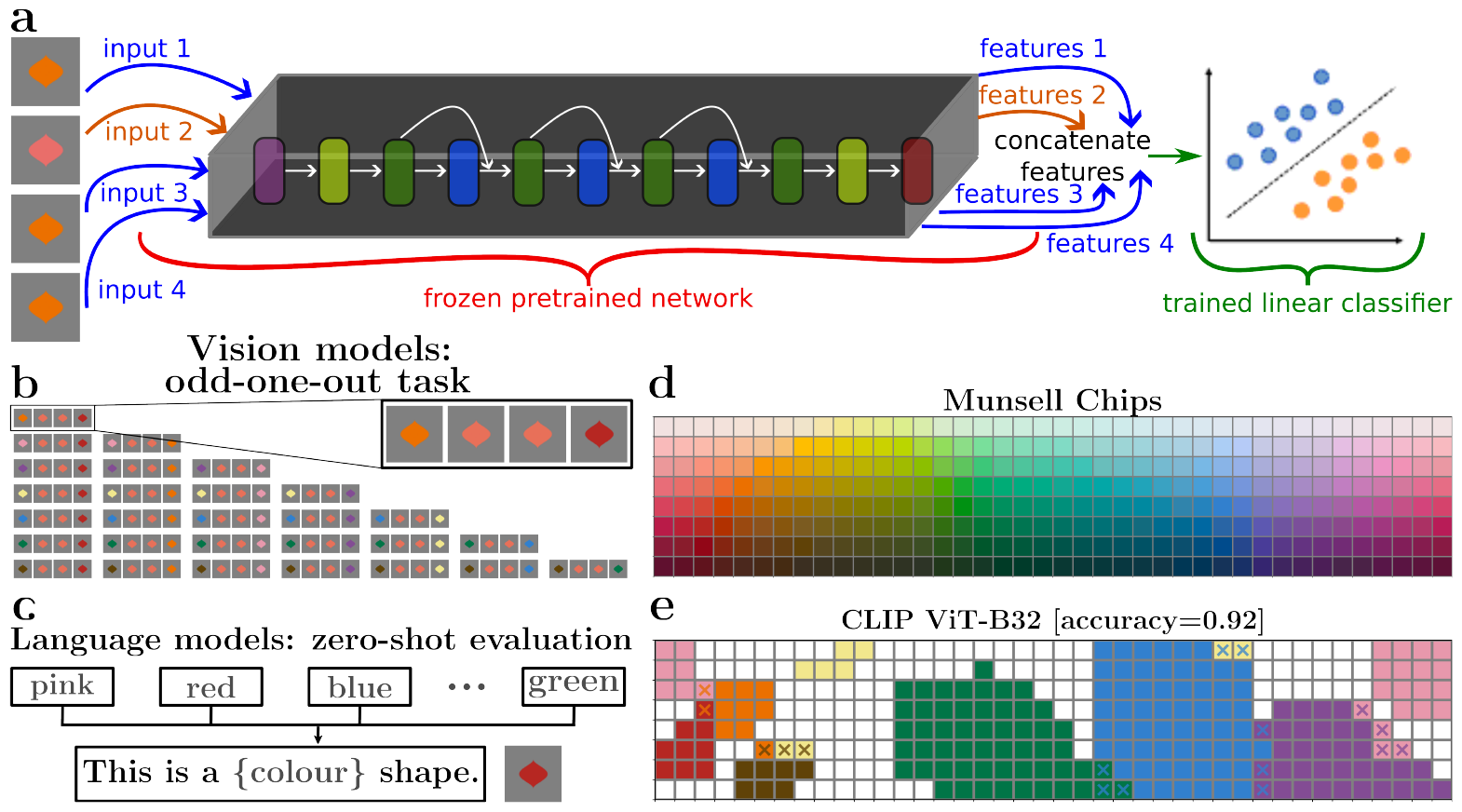
The Psychophysical Framework for Assessing Colour Categories in Artificial Neural Networks. Panel **a**: Linear classifier trained on features from a frozen pretrained network for a four-part oddone-out task. Panel **b**: Vision layer assessment using conflicting odd images—test colour presented alongside two focal colours, category determined by the non-selected focal colour, systematically repeated for all pairs of focal colours to eliminate bias. Panel **c**: Language model colour category testing through zero-shot evaluation. Network prompted with eight phrases, the category based on the term with the highest probability. Panel **d**: Displaying 320 Munsell chips as test colours. Panel **e**: Comparison of colour categories between one example network and human data [6, 41]. Filled cells represent network outputs, mismatches indicated by a cross coloured based on human data. White cells lack a unique colour category from the eight terms examined.

For the language layers of the CLIP models, we conducted direct psychophysical experiments without intermediary steps (see insert **c** in Fig. 1). Each Munsell chip underwent evaluation through eight phrases corresponding to different colour terms. We used the template “This is a {colour} shape.”, where “{colour}” is one of the eight colour terms. The network’s output, representing the probability score for each phrase matching the image, determined the final colour category. This process was repeated for all 2904 shapes, resulting in a total of 23,232 trials for each Munsell chip (2904*×*8).

Directly querying a pretrained vision model about colour categories is not feasible. To address this, we extracted features from a frozen pretrained network and trained a linear classifier for a four-part colour discrimination task (see insert **a** in Fig. 1). During testing, conflicting odd images were introduced to assess the categorical perception of vision models (see insert **b** in Fig. 1). The test colour (Munsell chip) was paired with two focal colours (e.g., orange and red). The network’s choice of the odd image determined the category of the test colour; for example, if the red focal colour is selected as the odd image, it indicates that the network grouped the test-Munsell chip into the orange colour category. Recognising the possibility that the test-Munsell chip could be neither red nor orange, we systematically tested it against all twenty-eight pairs of focal colours 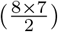. This procedure was repeated for all 2904 shapes, and the positions of focal colours were swapped to ensure unbiased results. In total, 162,624 trials were conducted for each Munsell chip (2904 *×* 2 *×* 28).

### 2.1 Role of language

In our investigation, we analysed four pretrained networks resulting from a combination of two tasks, CLIP (text-image matching) [31] and ImageNet (object recognition) [15], and two architectures: Vision Transformer (ViT-B32) [16] and Convolutional Network (ResNet50) [21]. We examined the networks at six different layers to elucidate the role of low-, mid-, and high-level visual representation in explaining categorical colour perception. Fig. 2 illustrates the accuracy of predicting human data, measured by assigning the same colour category for each Munsell chip. Our findings reveal that unimodal vision models can explain up to 76% of human data. In contrast, multimodal language-vision models achieve higher accuracy, reaching up to 95% with their language component and notably 89% even without the language component when exclusively testing the vision layers. These results underscore the dual role that language plays in categorical colour perception: a significant portion of human data is explained independently of language, while language-vision models show a 16% improvement in explaining human data, even when tested exclusively with their vision modality (similar to nonverbal psychophysics). Interestingly, testing with the language module (similar to verbal psychophysics) increases accuracy by a moderate 5%, suggesting that language shapes the development of colour categories, but the resulting representation is language-independent.

**Fig. 2.**
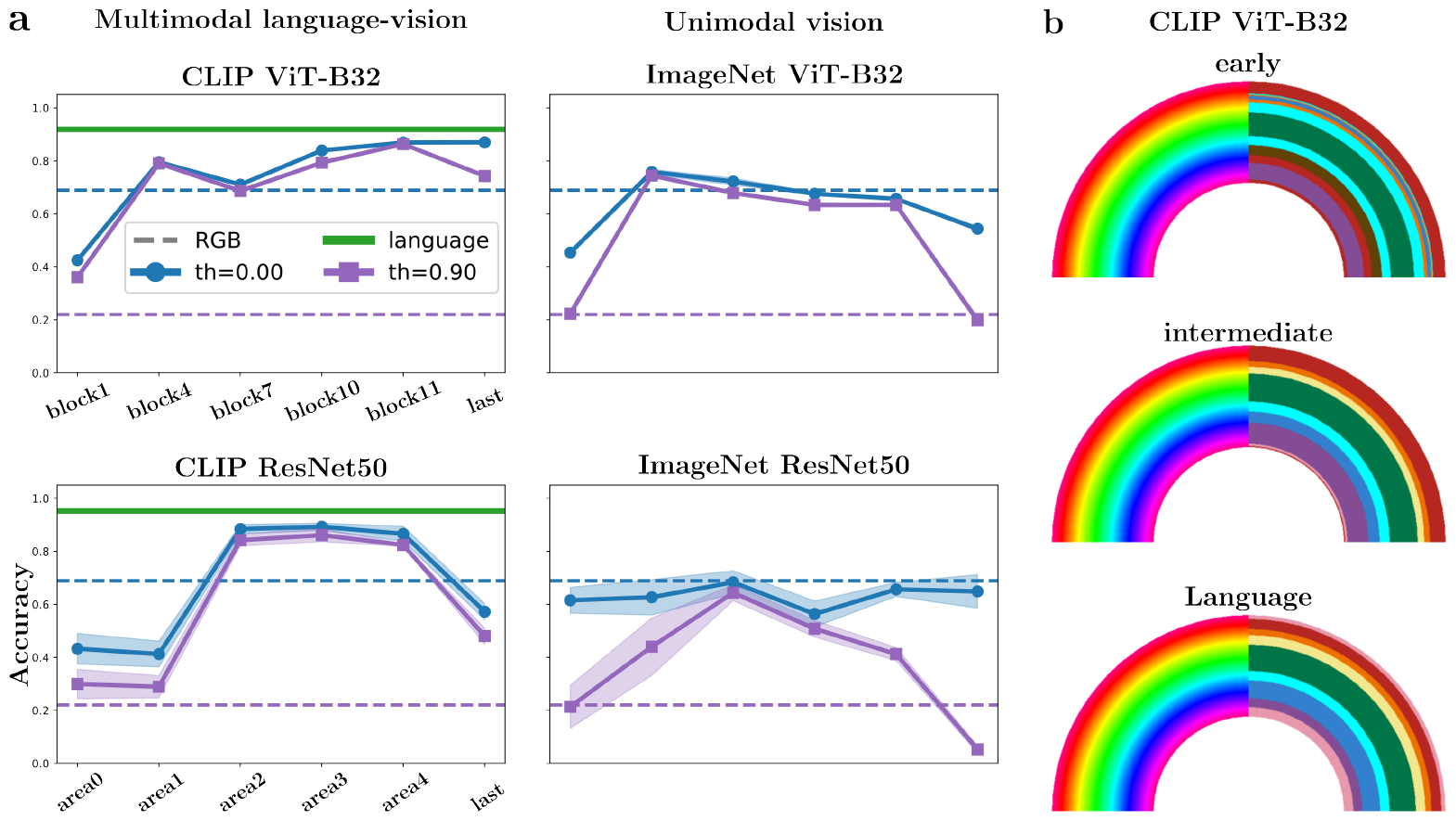
The Influence of Language on Colour Categories. Panel **a**: Shows accuracy in matching human data across six layers of four networks. Blue curves include all results, while purple curves indicate outcomes thresholded at 90% confidence. Transparent regions depict one standard deviation among five instances of linear classifiers trained with the same pretrained network (see Methods). The green horizontal line marks the accuracy when testing the network with the language module. Dashed horizontal lines represent colour categories based on Euclidean distance in RGB (networks’ input colour space). Panel **b**: Displays a rainbow image with continuous hue arches on the left. On the right, colour categories are obtained from an example network at three different layers.

To contextualise the accuracy of networks, we compared them to the RGB baseline. Given that the input colour space to networks is RGB, we defined a categorical model that computes the Euclidean distance to focal colours, with the smallest distance assigning the category of a Munsell chip. This baseline achieved a high accuracy of 68% in explaining human data, equivalent to the accuracy achieved by ImageNet ResNet50. However, when we applied a threshold to the results for higher confidence, the RGB accuracy substantially dropped to a third, whereas the accuracy of the networks did not change considerably (compare the purple and blue curves in Fig. 2). These results indicate that the input colour space is not the primary determinant of categorical colour perception.

Undertaking a qualitative analysis, the right panel of Fig. 2 presents the outcomes of a network prediction on a rainbow image. The arches of the rainbow, sharing identical values in saturation and value, display a continuous increase in hue by one degree. Despite the absence of any physical discontinuity in the rainbow arches, we distinctly perceive them in different colour bands. How do artificial networks interpret this image? In this experiment, we evaluated networks utilising nine colour terms, including the teal/turquoise category, given its qualitative visibility in the rainbow image and widespread usage [28]. Our observations reveal that the early layer differs significantly from our human colour perception, as it categorises bluish pixels as red and brown. In contrast, the intermediate representation closely mirrors how humans would categorise the rainbow image, except for the purple/pink split, almost entirely classified as purple. The language layer resolves this discrepancy by adjusting the purple and pink categories, perfectly aligning with our perception of the rainbow image.

### 2.2 Effect of visual task

The Taskonomy dataset [49] consists of twenty-four pretrained networks with an identical encoder architecture (ResNet50), trained on the same set of images for various visual tasks, spanning from low-level edge detection to mid-level depth estimation and high-level object classification. This dataset offers a unique opportunity to investigate the impact of a network’s functional role (the visual task a network is optimised towards) on its categorical colour representation. Employing the same analysis as detailed earlier, we scrutinised the networks at six different layers.

A significant disparity is evident among networks in predicting human data— assigning the same colour category for each Munsell chip (see the left panel of Fig. 3). The networks are ranked based on their peak accuracy across six layers, highlighting a substantial gap between the best-performing network, achieving 82% accuracy, and the least-performing one, attaining 16% accuracy. On one end of the spectrum, networks optimised for high-level semantic tasks, like “Object Classification”, consistently demonstrate human-like categorical representations. Conversely, networks performing 2D visual tasks, such as “2D Edge Detection”, consistently fall short of achieving human-like colour categories. Their predictive capability essentially hovers around chance levels across all layers, markedly lower than the baseline (Euclidean distance in RGB, the network’s input colour space). This implies that categorical colour representation is not a beneficial representation for networks trained on 2D visual tasks.

**Fig. 3.**
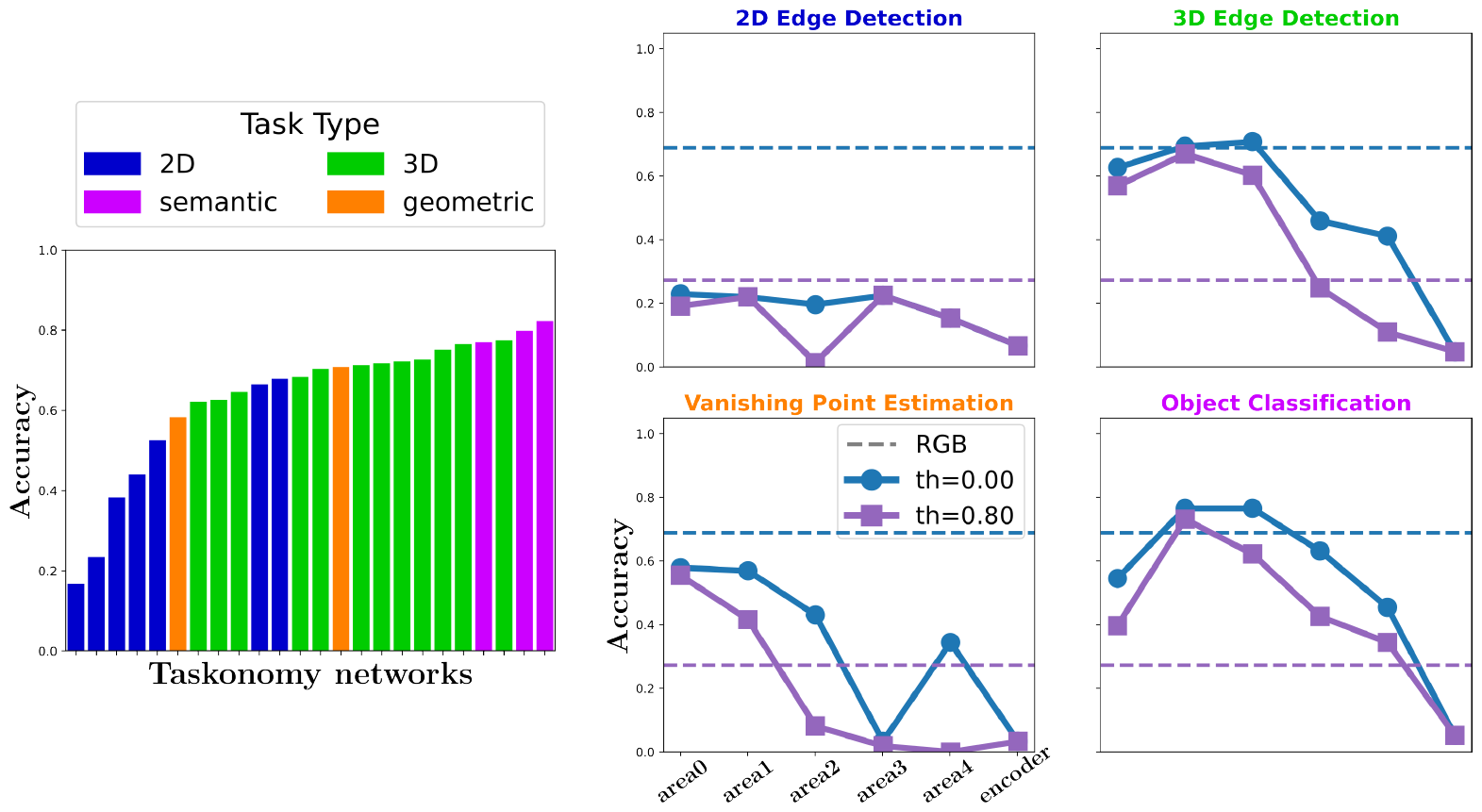
Effect of Visual Task on Colour Categories. **Left**: Ranks Taskonomy networks by their peak accuracy in explaining human colour categories. **Right**: shows accuracy in matching human data across six layers of four Taskonomy networks (see Fig. A3 for all twenty-four networks). Blue curves include all results, while purple curves indicate outcomes thresholded at 80% confidence. Dashed horizontal lines depict colour categories based on Euclidean distance in RGB (networks’ input colour space). Networks names are colour-coded by task types [17]: 2D, geometric, 3D, and semantic.

The taxonomy we adopted to classify these networks into four groups (2D, 3D, geometric, and semantic) relies on established criteria from prior literature, including methods such as representational similarity analysis (RSA) [18] and feature transfer learning [49]. Remarkably, our analysis yields similar clusters: along the spectrum of explaining human data, 2D tasks are situated on the left, 3D tasks in the middle, and semantic tasks on the right. This distinction holds true even for equivalent perceptual tasks in different dimensions. For example, the network trained on “3D Edge Detection” achieves human-like colour categories, whereas its corresponding 2D networks do not (as observed in the right panel of Fig. 3). This pattern extends to other corresponding 2D/3D tasks, such as keypoint detection. Collectively, these findings suggest that the nature of the visual tasks a system is designed to perform strongly influences its representation of colour categories. It can be hypothesised that our categorical colour perception has evolved due to living in a three-dimensional space and tackling semantic tasks.

### 2.3 Internal representation

The comparison of networks/layers to human data has revealed a distinct division. Some networks/layers closely approximate human colour categories, while others fail to align with them. This raises the question of whether there is a fundamental difference in how these two groups of layers/networks represent colours. It is important to note that networks’ colour categories are determined through a winner-take-all operation on an eight-class distribution. This is essentially a discrete procedure where one colour wins the category while the rest are silenced. However, before the discretisation stage, the underlying representation is a continuous distribution of the winning ratio among pairs of colour categories, which is a matrix of size 8 *×* 8 (refer to Fig. A2 in the supplementary material). To compare the internal representations of colour categories in networks/layers, we calculated the average Spearman correlation coefficients on this eight-class confusion matrix for each Munsell chip.

The left insert in Fig. 4 presents a pairwise comparison of all probed layers in CLIP and ImageNet networks. Notably, the continuous representation (upper triangle) exhibits better agreement across networks/layers compared to the discrete categories (lower triangle). The average correlation in categorical distributions (continuous) across all layers of CLIP/ImageNet ViT-B32/ResNet50 networks is 0.63 *±* 0.12. In contrast, the percentage of matching colour categories (discrete) shows both a lower average and higher standard deviation (0.54 *±* 0.18), indicating that the underlying continuous representations are significantly more similar than the discrete colour categories. This heightened correlation in the underlying continuous representation is particularly evident within the layers of a single network (depicted by dark-bordered squares; *r*_*s*_ = 0.72 *±* 0.04), and it is notably pronounced in ViT networks.

**Fig. 4.**
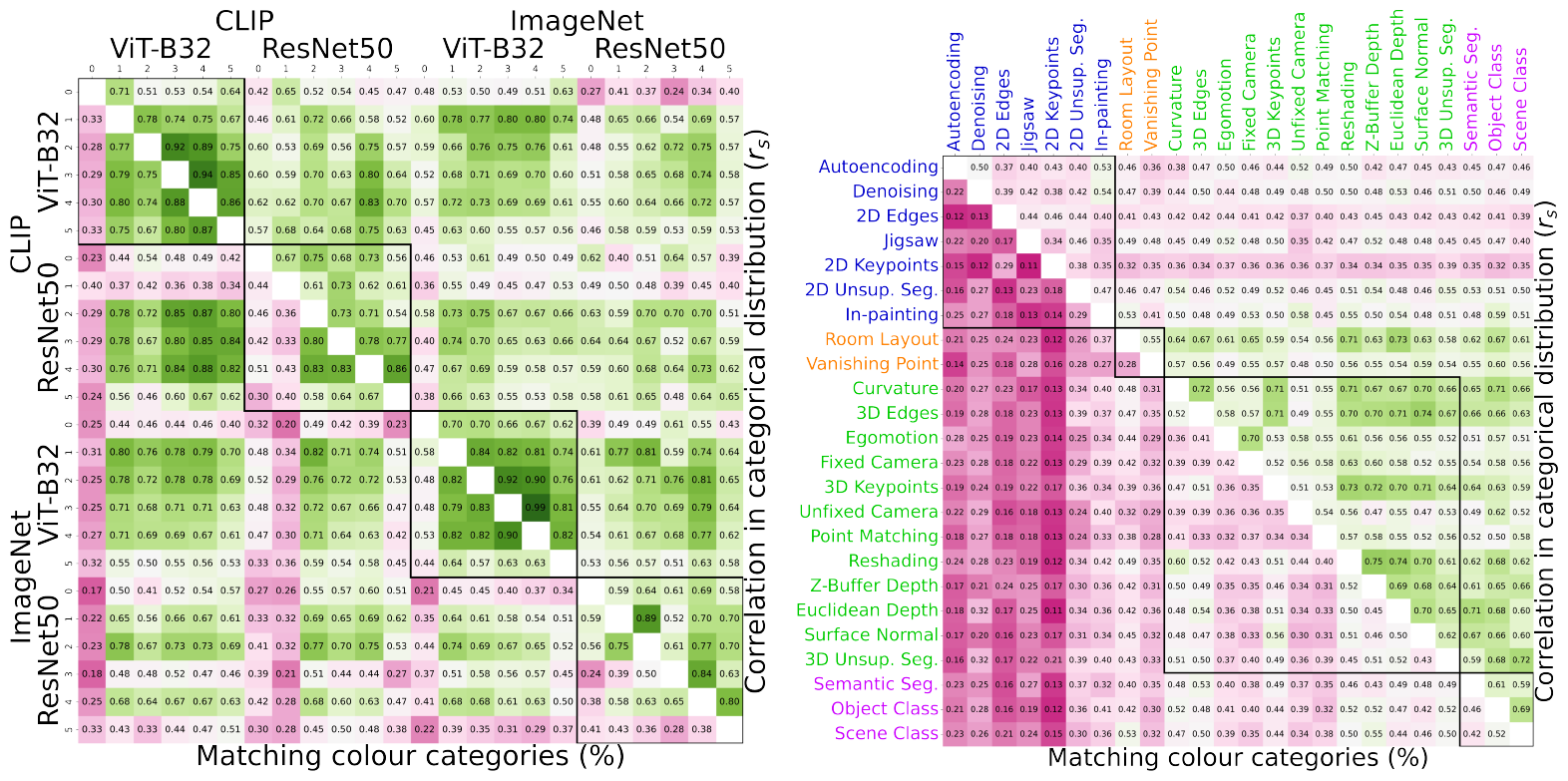
Comparison of Continuous and Discrete Representations. On the **left**, the upper triangular cells present Spearman correlations in the categorical distribution between pairs of layers, while the lower triangle indicates the percentage of matching colour categories. The dark-bordered squares represent layers within a single network. Cells are colour-coded, with green indicating 1 and purple indicating 0. On the **right**, the same format is applied to Taskonomy networks. The values in each cell are averages across the corresponding six layers in the networks. Network names are colour-coded based on task types [17]: 2D, geometric, 3D, and semantic. The dark-bordered squares delineate networks within a specific task type.

The right insert in Fig. 4 illustrates a parallel analysis conducted for the Taskonomy networks. The presented comparisons between networks are averaged over layerwise values (i.e., six layers). The first notable observation is the low ratio of matching colour categories across all 24 Taskonomy networks (purple cells in the lower triangle). This observation is not surprising, given the substantial variation in accuracy when explaining human data across different tasks (see Fig. 3 in the main text and Fig. A3 in the supplementary material). A second noteworthy pattern is the moderate green cells in the upper triangle, indicating a decent correlation (*r*_*s*_ = 0.65) in the categorical distribution of most visual tasks, except the networks trained on 2D tasks (blue labels). This strongly suggests that although the winner colour categories for these networks are notably different, the underlying representation is not significantly dissimilar.

We further scrutinised the networks’ continuous representation in relation to human colour naming consistencies data for British and German adults [46]. The language layers in CLIP networks exhibited a high correlation coefficient score (*r*_*s*_ = 0.65) aligning closely with the correlation between British and German speakers (*r*_*s*_ = 0.67). The vision layers in multimodal language-vision networks (CLIP) showed a similar correlation coefficient (maximum *r*_*s*_ = 0.63), while unimodal vision networks (ImageNet) showed a significantly lower correlation (*r*_*s*_ = 0.35). In fact, a strong correlation (*r*_*s*_ = 0.86) emerges between similarity to human colour naming consistency and accuracy in matching human colour categories. This comprehensive analysis suggests a meaningful relationship between networks’ continuous representation, human colour naming consistencies, and accuracy in replicating human colour categories.

**Fig. A1.**
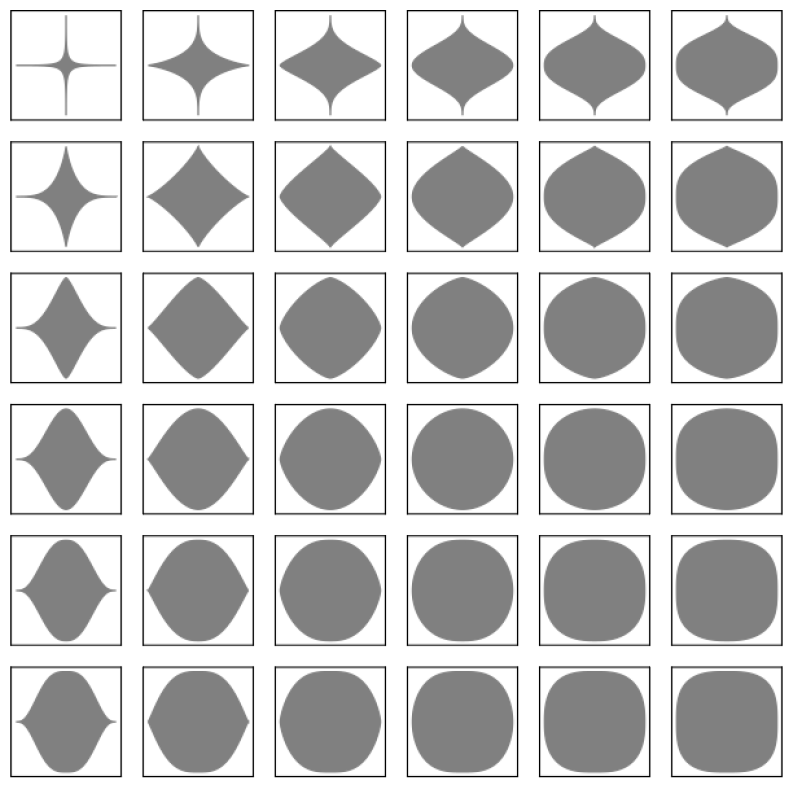
Example of thirty-six superellipse shapes obtained by keeping *a* = *b* = 0.5 and systematically varying the *m* and *n* values in Eq. A1.

## 3 Discussion

Communication plays an integral role in categorical colour perception, evident in our frequent use of colour names, even during inner speech. Recent studies, supporting the universalists’ standpoint, propose that efficient communication underlies the formation of colour categories [10, 42, 50]. This concept is also seen in the animal kingdom, where colour categories are intertwined with nonverbal communication needs like sexual mating [9]. Even in nonverbal human experiments indicating the emergence of colour categories independent of language, such as those involving stroke patients [39] and prelinguistic infants [40], the correlation between language and vision is inseparable, due to the nature of our brain. In the realm of artificial agents, this inherent language-vision correlation can be eliminated, allowing for models without language and communication components. This advantage has been leveraged in artificial neural networks for object recognition, revealing that human-like colour categories emerge to a considerable extent based solely on their utility for a particular vision task [14]. Our results advance this understanding by quantifying the contributions of each essential component—visual signals and linguistic factors. Notably, we find that a significant portion (about 80%) of human colour categories emerge in unimodal vision models. Nevertheless, a small yet important portion (about 20%) remains unexplainable purely on the basis of visual signals, which is clarified by the inclusion of multimodal language-vision models, underscoring the intricate interplay of these components in the development of categorical colour perception.

The utility of colour naming in communication is evident, as it is unfeasible to reference every discriminable tristimulus value with a unique colour name [26]. Hence, using distinct colour names for a broader range of hues proves efficient. However, the direct relevance of colour categories to a visual system is less apparent in the absence of communication or language interactions. To explore this, we examined Taskonomy networks, encompassing twenty-four distinct functional roles (i.e., visual tasks defining the optimisation loss) using an identical neural circuitry (i.e., ResNet50 encoder architecture) and training environment (i.e., exposed to the same set of images). The results resonate with the idea that the primary function of colour is to provide information relevant to behavioural tasks in the natural environment [11] by revealing the task-dependent nature of colour categories [44, 25] in a dualistic manner. While human-like categorical colour representation does not emerge in networks trained on 2D tasks, it is not scarce in other functional roles. This challenges the proposition of a unique connection between object recognition and colour categorisation [14]. Indeed, our findings suggest that, besides semantic tasks, 3D tasks such as shade parametrisation, depth estimation, and 3D edge detection yield human-like colour categories. The exact benefits of colour categories for specific functional roles, as opposed to others, remain to be investigated. However, categorical colour representation might be associated with foreground-background segmentation [20], a fundamental task continuously performed by infants in their daily lives, potentially explaining the early development of categorical colour perception in prelinguistic infants [27].

The first and second stages of colour processing, involving cone activation to different wavelengths of light and the antagonistic combination into colour opponency, are well-established [19] and reported to manifest in artificial neural networks [32, 3]. While these low-level mechanisms account for colour discrimination thresholds, they prove insufficient in explaining colour categories [37, 47]. Our experiments affirm this limitation; irrespective of the network’s architecture, modality, or training dataset, the initial layer does not exhibit any categorical effect. It has been postulated that, given the inadequacy of low-level mechanisms in elucidating colour categories, higher-level cognitive processes influenced by linguistic terms mediate categorical colour perception [37]. Our results challenge this notion by demonstrating that beneath different colour categories, a similar continuous colour representation may exist. This observation is independent of language modulation and consistently emerges in unimodal vision models. The involvement of high-level visual processes in categorical colour encoding remains uncertain [12, 7, 48]. However, our findings do not support this perspective in artificial networks, as the peak accuracy in matching human colour categories is never observed in the final layer. Conceptually, this aligns with the idea that high-level concepts should not strongly associate their representation with colour categories (e.g., recognising an apple based on its shape rather than its colour), and low-level processes should favour generic features in a continuous colour representation (e.g., detecting edges based on fine details of pixel values rather than coarser colour categories).

The connection between continuous colour perception and discrete colour categories remains a major challenge in the field of colour science [45, 38]. We posit that a meticulous analysis of intermediate layers in artificial networks can offer valuable insights into this intricate issue. In our experiments, Taskonomy networks (ResNet50 architecture) consistently show categorical colour representation emerging early in area 1, with peak accuracy sustained at mid-level representation (usually areas 1-2), followed by a rapid decline in deeper layers. Similar patterns are observed in ImageNet and CLIP networks (across both ResNet50 and ViT-B32 architectures). However, language models experience a more moderate drop in deeper layers, likely attributed to language modulation interacting directly with the final visual layer. These findings suggest that categorical colour representation is a mid-level feature in artificial neural networks, loosely aligning with the observation in rhesus monkeys that mechanisms for encoding colour categorically should occur earlier than visual area V4 [43]. The investigation into why mid-level mechanisms favour a categorical colour representation remains a subject for future exploration, yet insights from artificial neural networks propose that they may hold the key to advancing our understanding of categorical colour perception [30].

## 4 Methods

All research materials, including the source code for training/testing artificial neural networks and analysing the data, are openly accessible on our GitHub project page: https://arashakbarinia.github.io/projects/colourcats/.

### 4.1 Stimuli

The stimulus consists of uniformly coloured foreground and background images (see Fig. 1), offering the flexibility to dynamically adjust their surface colours for testing each Munsell chip. Foreground shapes are systematically selected from a set of 2904 geometrical shapes (refer to Appendix A.1 for details). The images are sized at 224 *×* 224 pixels, consistent with the image resolution utilised during the pretrained stage of all the examined networks.

We compared the colour categories of the networks to the human data from [6, 41]. The reported accuracies are the average over the union of ground-truths provided by these two studies, which encompassed 209 Munsell chips.

## 4.2 Pretrained networks

We investigated twenty-eight artificial neural networks trained on three distinct datasets:

- **ImageNet** [15]: containing 1.5 million images spanning over 1000 object categories. We investigated two architectures, namely ResNet50 [21] (a convolutional network) and ViT-B32 [16] (a transformer network). The pretrained network weights for both architectures were obtained from *torchvision*^1^.
- **CLIP** (Contrastive Language-Image Pretraining) [31]: comprising multimodal language-vision networks. These models contain a transformer text encoder and an image encoder that are jointly optimised to predict correct pairings of image-text batches. Our exploration involved two types of image encoders within the CLIP framework, namely CLIP ResNet50 and CLIP ViT-B32.
- **Taskonomy** [49]: encompassing around four million images, predominantly depicting indoor scenes, with labels for 24 computer vision tasks. The dataset provides pretrained weights of an encoder-decoder for all visual tasks. We focused our investigation on the encoder modules, which maintain an identical ResNet50 architecture across all tasks.

## 4.3 Colour-discriminator linear classifier

We applied the linear probing technique [5] to evaluate the categorical representation of colours in unimodal vision networks. This method enables the execution of psychophysical experiments with artificial neural networks, employing paradigms similar to human studies [4]. Furthermore, it permits the extraction of features at any layer, thereby providing a means to investigate intermediate features. The implementation utilised the osculari Python package [2] for a four-part odd-one-out colour discrimination task (see insert **a** in Fig. 1). Throughout training, four images were individually input into a frozen pretrained network (i.e., unaltered weights). Extracted features were then concatenated into a single vector and fed into a linear classifier. This classifier was trained to distinguish the odd image, identical to the other three in all aspects except for its foreground colour. To eliminate colour bias in the linear classifier, foreground colours were randomly selected from a uniform RGB distribution, while the background was uniformly chosen from achromatic colours (i.e., *R* = *G* = *B*). Stochastic gradient descent (SGD) optimised the linear classifier over 150,000 iterations.

For each architecture, we assessed colour categories at six distinct layers, comprising five intermediate layers and the final layer. In ResNet50, the intermediate layers are defined as areas 0 to 4, while in ViT-B32, they correspond to blocks 1, 4, 7, 10, and 11. Although we endeavoured to align the intermediate layers across architectures by selecting layers at similar depths, it is important to note that an exact match is unattainable due to the inherent differences in their architectures.

To bolster the robustness of our findings, we trained five instances of the colour-discriminator linear classifier, utilising the identical features extracted from the pretrained networks. The resulting colour categories from these five instances exhibit remarkable consistency (refer to the almost imperceptible standard deviations in insert **a** of Fig. 2). This observation strongly implies that the colour categories assigned by artificial networks are predominantly shaped by features acquired during their pretraining phase, with minimal influence from the colour-discriminator linear classifier.

During testing, we assessed the categorical characteristics of pretrained networks by introducing conflicting odd images (see insert **b** in Fig. 1). In this scenario, the background colour is always mid-grey (i.e., *R* = *G* = *B* = 128). Two of the four images are identical, featuring the test colour in their foreground, while the other two images display the focal colour of two distinct categories in their foregrounds. The unselected focal colour indicates the colour category of the test colour from the perspective of the network. To mitigate bias associated with our categorical colour perception, this procedure is repeated for all twenty-eight pairs of colour categories 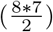. This procedure was repeated for all 2904 shapes, and the positions of focal colours were swapped to ensure unbiased results. In total, 162,624 trials were conducted for each Munsell chip (2904 *×* 2 *×* 28).

## Acknowledgements

This research received funding from Deutsche Forschungsgemeinschaft SFB/TRR 135 (grant number 222641018) TP S. An abstract of this work was presented at the European Conference on Visual Perception 2023 [1]. We express our gratitude to Christoph Witzel for supplying the human colour naming consistency data.

## Appendix A Extended data

### A.1 Stimuli shapes

To create the test shapes in our study, we employed the superellipse, defined in the Cartesian coordinate system as the set of all points (*x, y*) satisfying the equation

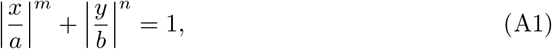

where *a, b, m* and *n* are positive numbers. Fig. A1 depicts thirty-six examples of these superellipse shapes. The selection of a systematic geometrical shape serves the purpose of exploring the interaction between object shape and colour perception, although this aspect falls outside the scope of the current article.

### A.2 Raw experimental data

The exhaustive examinations conducted to evaluate the categorical representation of colours in vision layers through linear probing yield an 8 *×* 8 multi-class confusion matrix, as illustrated in Fig. A2. Several noteworthy aspects of this matrix warrant attention:

- Higher values indicate a robust category effect, while values close to 0.5 (chance level) suggest an absence of categorical representation.
- The summation of winning ratios for a specific pair of colours may not necessarily equate to 1.0. For example, in Fig. A2, the sum of winning ratios for the orange-red colour categories is 0.99. The remaining percentage pertains to scenarios where the test colour has been selected as the odd image. This can be construed as noise in the linear classifier. Overall, the magnitude of this noise is minimal, accounting for only 0.02 across all layers.
- The relationship between colour categories is not entirely transitive. In Fig. A2, although orange prevails over red 78% of the time, when compared to brown and purple categories, red obtains a marginally higher winning ratio (1% more, i.e., 100 versus 99). Whether this discrepancy is attributable to noise in the linear classifier or signifies the non-transitive nature of colour categories remains unclear. Nevertheless, similar to the aforementioned point, the impact is exceedingly marginal.

**Fig. A2.**
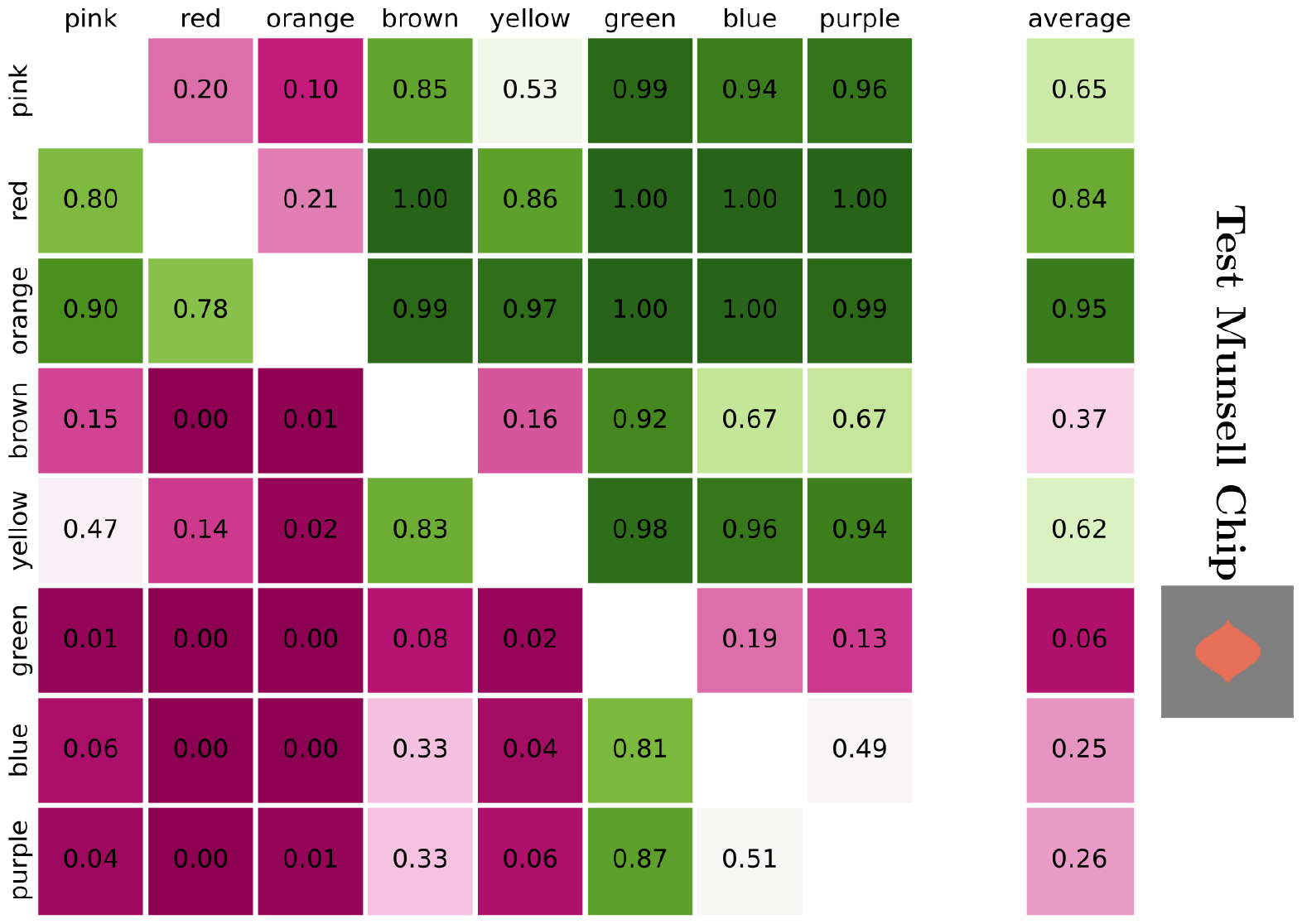
Distribution of Winning Colour Categories (Derived from Block-10 of CLIP Vit-B32). Each cell denotes the percentage of a category selected as the colour for the illustrated test-Munsell chip. The values in the upper and lower triangles may not necessarily add up to 1; the remaining percentage (typically minimal) indicates instances where neither category is chosen. The numbers reflect the average across 5810 tests. Cells are colour-coded, with green representing 1 and purple representing 0.

### A.3 Taskonomy results

Fig. A3 illustrates the accuracy in matching with human colour categories for all twenty-four Taskonomy networks across six layers. The networks are arranged in ascending order based on their peak accuracy in explaining human data. Notably, in the top two rows, all networks grouped under the 2D task type [17] demonstrate inferior performance compared to the RGB baseline. This observation implies that categorical colour representation is inconsequential to their functional role.

**Fig. A3.**
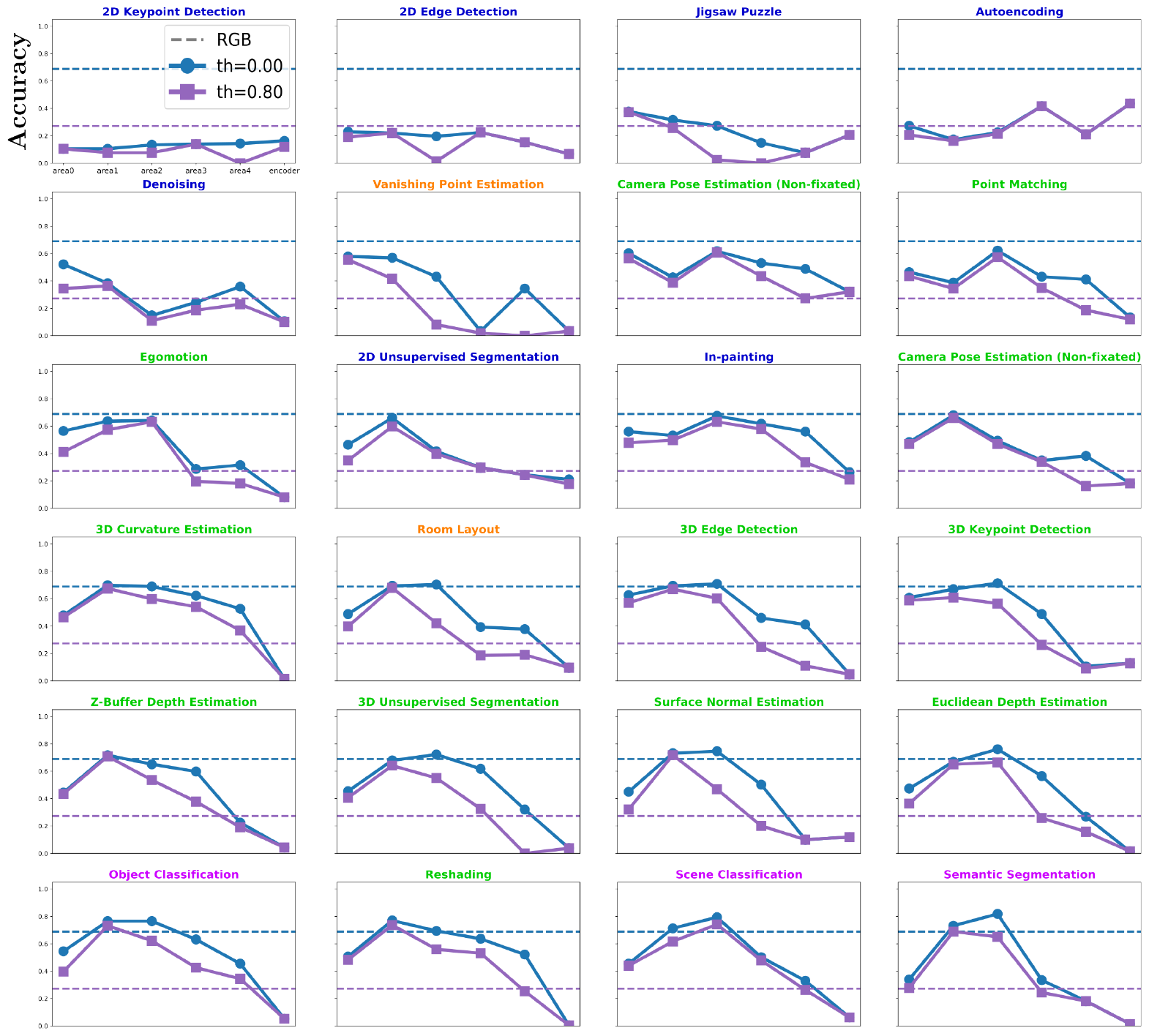
Results of Taskonomy Networks. Accuracy matching human data in six layers of 24 Taskonomy networks. Blue curves include all results, while purple curves indicate outcomes thresholded at 80% confidence. Dashed horizontal lines depict colour categories based on Euclidean distance in RGB (networks’ input colour space). Networks names are colour-coded by task types [17]: 2D, geometric, 3D, and semantic.

## Footnotes

1 https://pytorch.org/vision/stable/models.html

